# Exploring the Exclusive Isolation of *Pseudomonas syringae* in *Peltigera* Lichens via metabolite analysis and growth assays

**DOI:** 10.1101/2024.09.06.611675

**Authors:** Natalia Ramírez, Diana Vinchira-Villarraga, Mojgan Rabiey, Margrét Auður Sigurbjörnsdóttir, Starri Heidmarsson, Oddur Vilhelmsson, Robert W. Jackson

## Abstract

The specific association of the potentially plant-pathogenic *Pseudomonas syringae* with *Peltigera* lichens raises questions about the factors driving this host specificity. To explore this, the metabolic profile of seven lichen species belonging to three genera (*Cladonia, Peltigera*, and *Stereocaulon*) was analysed using LC-MSMS. Additionally, we assessed the growth of *P. syringae* strains in media supplemented with extracts from each lichen species. This revealed that *Peltigera* exhibits lower metabolite richness compared to other genera, but shows a higher chemical investment in specific compounds. Growth kinetics showed comparable *P. syringae* growth across lichen-supplemented media, except for *C. arbuscula* and *Cladonia* sp., where the former exhibited lower growth rates. Inhibition assays with lichen extracts showed no inhibition of *P. syringae*. The lichen metabolome is predominantly composed of lipids and organic acids. Furthermore, specific compounds, such as aminoglycosides, may facilitate *P. syringae* presence in *Peltigera* by inhibiting *Bacillus subtilis* and other antagonists. Additionally, compounds absent in *Peltigera*, like anthracene, might serve as a carbon source inhibitors like *B. velezensis*.

## Introduction

Lichens are a widespread and diverse group of cryptogam species formed by a symbiosis between fungi and photosynthetic symbionts [1,2]. The lichen thallus comprises intertwined fungal hyphae and photosynthetic cells, consisting of either algae, cyanobacteria, or a combination of both. Beyond these constituents, lichens also host internal communities of bacteria [3,4], fungi [5,6], archaea [7,8], and viruses [9]. Lichens rely heavily on atmosphere water and their microbiome, which plays a crucial role in their survival in harsh environment [3,10]

Given their widespread distribution, lichens can be found in environments with variable climates, including subpolar oceanic zones such as Iceland. Terricolous lichens are a significant constituent of Iceland’s vegetation, accounting for approximately 3.64% of vegetation coverage [11]. Some of the most common Icelandic lichens include members of the genera *Peltigera, Stereocaulon*, and *Cladonia. Stereocaulon* is a cosmopolitan lichen genus comprising approximately 125 fruticose species [12], including the three-part symbiotic species *S. alpinum* and *S. vesuvianum* [13]. *Cladonia* is a genus characterised by a primary crustose or squamulose thallus followed by a secondary fruticose thallus. This genus encloses approximately 475 species [14], from which *C. arbuscula* and *C. chlorophaea* adhere to a bipartite arrangement involving an algal partner [10,15]. The lichen genus *Peltigera* is typically foliose and exhibit broad lobes [16]. This diverse genus encompasses around 100 lichen species [17], with examples like *P. leucophlebia* and *P. apthosa* containing a combination of both algae and cyanobacteria as photobionts [18], and species such as *P. membranacea* that exclusively harbour cyanobacteria [19].

Due to their complex nature, lichens also possess a broad metabolism, drawing considerable interest due to their distinct chemistry and biotechnological prospects. Lichen’s secondary metabolites are intricate yet non-essential small molecules that can comprise 0.1% to 30% of the lichen thallus’ dry weight [20]. Throughout history, the compounds derived from lichens have been thought to be synthesised by the dominant fungal partner. However, recent studies indicate that photobionts and bacteria associated with lichens also contribute to a variety of potentially valuable molecules, and although some of those metabolites may not directly benefit the microorganisms, they are advantageous to the overall lichen [21].

Over 1050 recognised lichen metabolites are catalogued in the Lichen DataBase (LDB). These metabolites have multifaceted functions within the lichen symbiosis and their ecological niche. Some of them serve as allelopathic agents, influencing the growth and survival of neighbouring organisms, thereby giving lichens a competitive advantage [22]. Others affect the palatability of lichens to herbivores, acting as a form of defence. Furthermore, certain secondary metabolites enhance the permeability of the cell membranes of phycobionts, the algal partners within lichens, potentially aiding in nutrient exchange and survival in challenging conditions. Additionally, these metabolites serve as shields against excessive ultraviolet (UV)-B radiation, helping to protect the photosynthetic components of lichens from harmful UV rays [22]. Moreover, it is noteworthy that a substantial portion of studied lichen species synthesises substances with varying degrees of antimicrobial activity [23], which could have implications for their interactions with other microorganisms in their environment. Overall, these protective compounds collectively contribute to the unique and enduring conditions for lichen existence, resulting in remarkable longevity, with certain lichens living for several millennia [24].

Earlier investigations in Icelandic lichens of the genera *Peltigera* indicated a particular prevalence of *P. syringae* as part of their microbiome [11,25]. Interestingly, *P. syringae* was isolated consistently from only this lichen genus out of ten evaluated. Furthermore, these bacteria were found in 4 different *Peltigera* species collected. The genetic lines of *P. syringae* discovered in the Icelandic region are remarkably monophyletic, suggesting independent evolutionary trajectories from *P. syringae* populations in other global regions. These monophyletic haplotypes are observed across various phylogroups (PGs) [25]. Notably, certain *P. syringae* strains associated with PG01 and PG02 underwent pathogenicity and fitness assessments across different plant species, revealing pathogenicity levels comparable to those of epidemic *P. syringae* strains such as DC3000 or CC0094 [11].

Following the exclusive isolation of *P. syringae* from lichens of the *Peltigera* genus, we became intrigued by the potential differences between *Peltigera* and other lichen genera. Initially, we hypothesised that the lack of *P. syringae* isolated in other lichen species might be due to the higher water content in *Peltigera*, which is associated with an observated increase in the amount of culturable bacteria [11]. A second hypothesis considered the lichens’ chemistry. We hypothezised that a metabolite(s) within *Peltigera* lichens might be enhancing *P. syringae*’s survival or reducing the growth of known *P. syringae* antagonistic bacterial strains such as other *Bacillus velezensis* [26]. Conversely, non-*Peltigera* lichen genera might contain metabolites inhibiting *P. syringae* growth or enhancing the population of antagonistic microbes on their thallus, which could reduce *P. syringae* population.

In this context, this research aimed to determine whether the exclusive isolation of *P. syringae* from the lichen genus *Peltigera* was influenced by the lichens’ chemical environment. To achive this, three different approaches were taken. First, we analysed the metabolic profiles of lichens from different species from the *Peltigera, Cladonia*, and *Stereocaulon* genera to evaluate if there were distinctions between the different genera. Next, the growth profile of three lichen-isolated *P. syringae* strains and an outgroup bacteria, *B. velezensis*, was obtained in a medium supplemented with lichen extract to determine if the growth of the strains was enhanced by the presence of the lichen metabolites. Finally, the antibacterial activity of the lichen extracts was tested against selected *P. syringae* strains isolated from the Icelandic lichens, as well as a control strain, the *P. syringae* antagonistic bacteria *B. velezensis*.

### Experimental Procedures

#### Sampling lichens

In March 2023, sample collection occurred in the southwestern region, specifically in the snow-free areas of Iceland in that season, including Heiðmörk forest, Öskjuhlíð hill, and the shores of Elliðaá in Árbæjarstífla. Following collection, all samples were cleaned of other organisms attached, such as moss or debris. Promptly after, all lichen thalli were flash-freezed in liquid nitrogen *in situ* and stored at -80°C. A total of 31 samples were collected, representing three distinct genera: *Cladonia, Peltigera*, and *Stereocaulon*, and seven specific species: *C. arbuscula, C. chlorophaea, P. aphthosa, P. leucophlebia, P. membranacea, S. alpinum*, and *S. vesuvianum*. Additional details regarding the sampling process, dates, and location are available in Table A1. All specimens underwent morphological analysis, and corresponding vouchers have been deposited at the Icelandic Institute of Natural History.

#### Tissue preparation

After lyophilisation during 48-72 hours at -52°C in LyoDry Benchtop Freeze Dryer (MechaTech Systems), samples were mashed until reaching a dust-like consistency with the help of a pestle. Samples of the same lichen species were pooled together in equal amounts and stored at -80°C for metabolite extraction as detailed below.

For the metabolite extraction, 80 milligrams of each lichen were placed into a 2 mL Eppendorf tube, and 500 µL of 100% methanol (LC/MS grade, Fisher Scientific UK Limited) were added. The samples were thoroughly mixed using a vortex mixer for one minute and sonicated for 20 minutes at room temperature, with an ultrasound frequency of 37 kHz and a power of 80 W (Fisherbrand, FB15051). After ultrasonication, the samples were centrifuged at room temperature for 10 minutes at 13,000 rpm, and the resulting supernatant was collected in a new Eppendorf tube. This process was repeated thrice, with fresh solvent added to each round. The obtained crude extract was dried using a vacuum concentrator (Eppendorf® Concentrator Plus) at 45°C. Afterwards, the total chemical investment (hereafter referred to as chemical investment) was calculated as reported by Salazar et al (2018) measured as the average thallus dry-mass percentage of secondary compounds.

#### HPLC-MS/MS analysis

Before analysis, the samples were reconstituted to a final concentration of 10 mg ml^-1^ in 50% acetonitrile (ACN, LC/MS grade, Fisher Scientific UK Limited) and filtered through a PTFE 0.22 µm filter (Fisherbrand™). A pooled sample (QC) was created by combining 10 µl from each extract to monitor the analysis performance and correct batch effects (if observed). Similarly, 50% ACN was employed as a blank control and included in the batch alongside the QC samples, to be analysed once every ten (randomised) lichen samples.

For HPLC-MS/MS analysis, 10 µl of each extract was injected into a Dionex UltiMate 3000 chromatographic system coupled to a Q-Exactive mass spectrometer (Thermo Fisher Scientific, Bremen, Germany). This analysis utilised a C18 column (Hypersil GOLD, 100X2.1 mm, particle size: 1.9 µm, Phenomenex, Torrance, USA). The mobile phase consisted of 95% H_2_O (LC/MS grade, Fisher Chemical™) with 0.1% formic acid (FA, Op tima™ LC/MS Grade, Fisher Chemical™) as solvent A, and 95% ACN with 0.1% FA as solvent B. The flow rate was set at 250 µl min^-1^.

The chromatographic program was configured as follows: 1% B from 0-1 min, a gradient increase from 1 to 99% B between 1-40 min, an isocratic phase at 99% B for one minute, followed by a one-minute return to 1% B and a 10 min re-equilibration phase at 1 % B. Data-dependent acquisition of MS/MS spectra was conducted in Electrospray Ionization (ESI) positive and negative modes. ESI parameters included 40 arbitrary units (AU) sheath gas flow, 10 AU auxiliary gas flow, and 3 AU sweep gas flow. The auxiliary gas temperature was set to 100 °C, the spray voltage was set to 3.9 kV in positive mode, and to 3 kV in negative mode. The inlet capillary was heated to 275 °C, and the S-lens level was adjusted to 50 V.

The MS scan range was set from 100 to 1800 Daltons, with a resolution of 140,000 and one micro-scan. For MS acquisition, the maximum ion injection time was 100 ms with an automatic gain control (AGC) target of 1 × 10^6^. Up to five MS/MS spectra per duty cycle were acquired at m/z 200 of 17,500. The maximum ion injection time for MS/MS scans was 50 ms with an AGC target of 1 × 10^5^ and a minimum AGC target of 8 × 10^3^.

The MS/MS precursor isolation window was set to m/z 4.0, and the normalised collision energy was stepped from 25% to 35% to 45% with z = 1 as the default charge state. Dynamic precursor exclusion was set to 10 s. For additional details regarding the parameters, programs, and analyses, please refer to Table A2.

### Data pre-processing and *in silico* compounds annotation

After the HPLC-MS/MS analysis, raw files underwent conversion into centroid mode (.mzML) using Msconvert [28]. Subsequently, Mzmine 3.2.8 was employed for feature finding, chromatogram deconvolution, alignment, gap filling, feature filtering, and ion identity (IIN), as detailed in Table A2 [29]. The results were exported in the form of feature abundance tables (.csv), MS/MS spectra files (.mgf), and ion identity networking results (.csv).

To enhance data quality, solvent-related features were removed using the data acquired from the solvent blank samples. Furthermore, features with a relative coefficient variation exceeding 30% in the QC samples were filtered out. In addition, adducts and in-source fragments identified by the IIN algorithm were eliminated to reduce the presence of overrepresented features, retaining only representative ions for subsequent analysis.

An extra spectral file (.mgf) was exported from MzMine for utilisation in SIRIUS 4.0, facilitating in silico compound classification through CANOPUS (CSI: FingerID; MSI, level 3) [30–32]. Feature class assignments were annotated using the ClassyFire chemical ontology output and the NPC library for specific cases where the annotation was more accurate and corresponded in chemical composition with the one from ClassyFire. Assignments with a probability exceeding 0.8 were considered valid, while those with a lower probability were categorised as “Unknown”.

### Chemical diversity assessment

To analyse the impact of the tested factors, genera, and species, a Permutational Multivariate Analysis of Variance (PERMANOVA) was conducted using the adonis2 function in the ‘vegan’ package (R 4.1.0). This involved assessing Bray-Curtis dissimilarity matrices created for each dataset. NMDS plots were generated using ggplot2 to visualise sample dispersion. Post-hoc pairwise PERMANOVA for each lichen genera was performed, adjusting for multiple comparisons with the pairwise.adonis function and Bonferroni correction.

For alpha diversity analysis of extracts, feature richness and the Simpson diversity index were calculated. Feature richness represented the number of detected features from each genera/species. The inverse Simpson index reflected the abundance-weighted diversity of metabolites in each sample. Alpha diversity indices differences were evaluated using a one-way ANOVA followed by a Tukey-Kramer HSD post hoc test (p<0.05).

Beta diversity was evaluated through principal component analysis (PCA), Hierarchical Clustering Analysis (HCA), and orthogonal partial least-squares discriminate analyses (OPLS-DA) in Metaboanalyst 6.0. OPLS-DA models were constructed to compare genera and species. Variable Importance in Projection (VIP) features with scores higher than 1.5 were selected as discriminant variables. The relative abundance of discriminant features was evaluated with unpaired t-tests and controlled for false discovery rates at 1%. Two-way ANOVA and Tukey’s multiple comparison tests were used to test differences between genera and/or species. PCA and OPLS-DA scores (R^2^X, R^2^Y, Q^2^Y) were employed to assess variance coverage and predictability, while permutation tests (n=999) were run and analysed to ensure the significance, stability, and non-randomness of OPLS-DA models.

### Bacterial growth assays

To evaluate the effect of the lichen metabolites on *P. syringae* growth, a kinetic growth curve of three *P. syringae* strains (EG201428, SU200101, HV200414) isolated from Icelandic lichens in a previous study [11], was performed using an M9-lichen supplemented media. As control, three non-lichen *P. syringae* isolates from PG2, PG10 and, PG13 (CC0094, CFBP1392, UB0246, and CC1557) and a non-*Pseudomonas* nor lichen-isolated strain, *B. velezensis* (KHWW02) were included. The bacterial strains were cultured overnight on King’s B [33] or Lysogeny broth (Composition per l, Tryptone 10g, Yeast extract 5g, NaCl 10g) at 27 °C with shaking (200 rpm). The bacterial suspensions were centrifuged and washed with phosphate-buffered saline (PBS, pH 7.4, Fisher Scientific UK Limited) three times to remove all KB or LB traces. After the washing, all strains were diluted in M9 without glucose until 0.4 Optical density 600 nm (OD_600_).

For this experiment, lichen crude extracts were obtained in 100% methanol as detailed previously, dried, dissolved in 20% methanol, and then added to Minimal medium (M9, without carbon source [34]) to reach a final concentration of 20 mg of extract per mL. Afterwards, the M9-lichen media was sterilised by filtration with a 0.22 µm PES filter and further diluted by mixing 100 µL of the M9-lichen broth with 100 µL of each bacterial suspension in a 96-well plate. This dilution allowed to reach a final extract concentration of 10 mg mL^-1^ and a bacterial OD_600_ of 0.2. The 96-well plate was placed into a TECAN SPARK® Multimode Microplate Reader (Tecan Trading AG, Switzerland) an incubated at 27 °C for 47 hrs with OD readings every 30 minutes. Three replicates of each treatment and duplicates per control (bacterial strains growing on M9 and KB) were included on each plate. The distribution of the plates was done by the combinations: *P. membranacea*-*P. apthosa*; *S. vesuvianum*-*S. alpinum*; *P. leucophlebia*-*C. chlorophaea* and *C. arbuscula*.

### Antibacterial activity assay of lichen extracts

The inhibitory effects of metabolites from 31 lichen samples, grouped by species, against various strains of *P. syringae* and *B. velezensis*, were assessed and interpreted using the Kirby-Bauer disk diffusion assay [35]. Briefly, 10 ml of KB broth [36]) was inoculated with 100 µl of the bacterial strains and incubated for 16 hours at 27 °C and 200 rpm. After incubation, the cell density was adjusted to 0.2 OD (OD_600_, equivalent to 10^8^ CFU ml^-1^), then further diluted 1:100 in 20 ml of KB [36]) agar (0.75% w/v agar) and poured into 90 mm Petri dishes.

Simultaneously, 25 µl of each lichen extract (10 mg mL^-1^ in 50% ACN) was added to a sterile 6 mm paper disk and allowed to dry for one hour. Once the solvent evaporated, three paper disks per extract were placed in each Petri dish and incubated for 24 hours at 27 °C. The formation of clear haloes surrounding the paper disk with no evident bacterial growth were considered as positive antibacterial activity. The diameter of the observed inhibition zones was recorded, and differences in antibacterial activity among treatments were assessed through one-way ANOVA, followed by a Tukey-Kramer HSD test (p < 0.05).

## Results and discussion

### *Peltigera* shows higher chemical investment focused on fewer compounds

To evaluate if the lichens from the genus *Peltigera, Cladonia*, and *Stereocaulon* had distinctive metabolic profiles, an untargeted metabolomics analysis was employed to obtain chemical profiles from eight lichen species (three *Peltigera*, three *Cladonia*, and two *Stereocaulon*). This analysis was conducted using LC-MS in ESI positive and negative modes, generating two datasets with 6527 features and 1504, respectively (Table A6). Multivariate data analysis was applied to statistically assess these features, revealing differences among the studied species. Additionally, metabolite annotation using GNPS and SIRIUS platforms provided details into specific compound presence. Out of the dataset (Table A6), 4.5% and 1.7% (corresponding to 297 and 25 features) were annotated (MSI level 3) in the positive and negative modes, respectively.

The *Peltigera* lichens demonstrated a higher chemical investment (extract weight/lichen dry weight ratio), ranging from 12% to 13%, in contrast to the considerably lower investment observed in the *Cladonia* and *Stereocaulon* lichens (2-4%). However, non-*Peltigera* lichens exhibited higher richness (number of detected features) than *Peltigera* in both polarities (positive – average richness in *Peltigera, Cladonia*, and *Stereocaulon*: 717, 1084.78, 1086.17, respectively; negative – 45.44, 81.72, and 96.75) (Figure 1). *Peltigera* shows a lower number of metabolites in both polarities, however in the positive mode there are no differences in the diversity based on Simpson reciprocal index test, while in the negative mode dataset, there were differences among all genera with *Peltigera* having the lowest score and *Stereocaulon* the highest (p <0.05). In terms of metabolite evenness based on the Simpson index, *Peltigera* samples showed significantly higher levels compared with *Cladonia* and *Stereocaulon* (*Cladonia*-*Peltigera* p= 0.0002; *Stereocaulon*-*Peltigera*, p= 2.044 × 10^−5^), while *Cladonia*-*Stereocaulon* comparison indicated a more balanced abundance of the compounds isolated from *Peltigera* samples (p= 0.4563). While in the negative polarity, *Stereocaulon* exhibited higher evenness compared to the other two genera, which did not differ (*Peltigera*-*Stereocaulon*, p= 1.14 × 10^−11^; *Cladonia*-*Stereocaulon*, p= 2.53 × 10^−14^; *Peltigera-Cladonia*, p= 0.4665). This suggests that *Peltigera*’s chemical composition is characterised by a substantial concentration of a few metabolites, while other lichen species display a more even distribution of a wider variety of metabolites in relatively similar quantities. The substantial allocation of energy and carbon resources towards the production of these secondary metabolites suggests they may have significant physiological and ecological roles [37].

**Figure 1.**
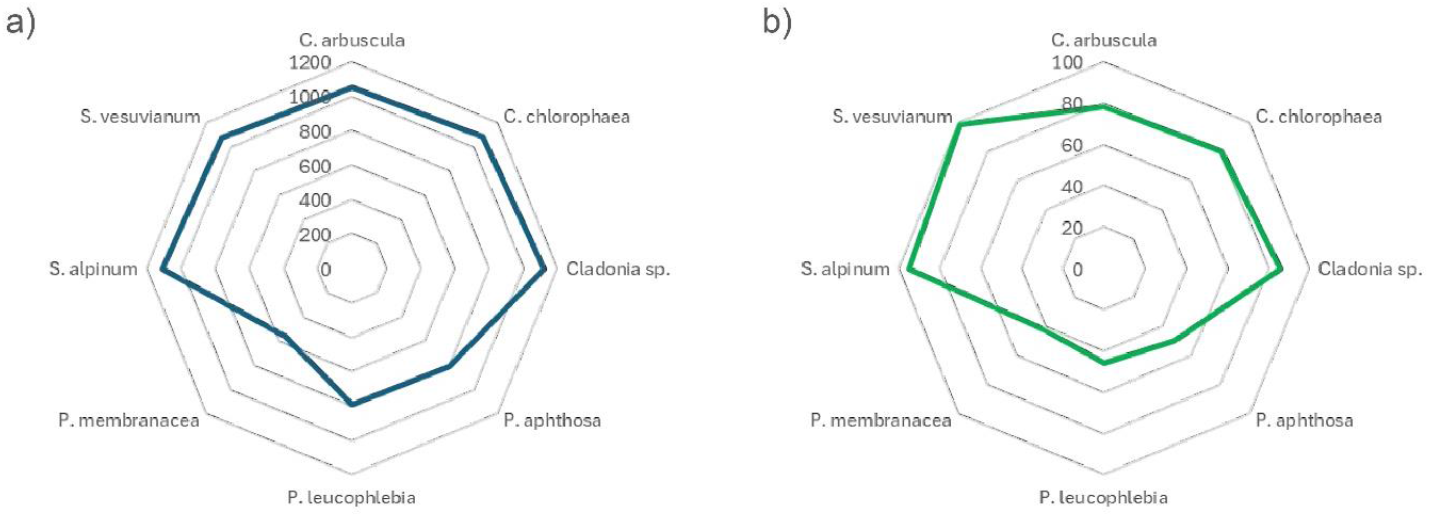
Average richness of metabolites within lichens differs across species. A) Positive Electrospray Ionization (ESI) metabolites and b) Negative ESI metabolites. The blue and green lines show the number of features detected. The graph was created by package fmsb in R studio.

Since *P. syringae* strains were isolated from all *Peltigera* species sampled, we analysed the *Peltigera* chemical profiles for three species of *Peltigera* to determine their common metabolites and evaluate if there could be a shared pool of metabolites that could support *P. syringae* colonisation within the *Peltigera* microbiome. Notably, the datasets obtained showed that all the evaluated species have a distinctive metabolic profile. *P. membranacea* showed less diverse metabolic profile with 2060 features detected, while in *P. leucophlebia* and *P. aphthosa* 2939 and 3005 features were observed, respectively (Figure 1) (Richness difference *P. membranacea-P. leucophlebia*: 5.38 × 10^−9^; *P. membranacea-P. aphthosa*: 1.70 × 10^−8^, and *P. leucophlebia-P. aphthosa*: 6.30 × 10^−1^). Despite this, there is a slightly higher chemical investment in *P. membranacea*, suggesting that this species may concentrate its metabolic production on a smaller number of compounds, as described before.

Regardless of their differences, there was a shared core metabolome in the three evaluated species constituted by 481 features that did not significantly vary in terms of abundance (p>0.05). These features belonged mainly to the classes prenol lipids, glycerophospholipids, carboxylic acids and derivatives, and fatty acyls chemical families, being grouped specifically in the glycerophosphocholines, amino acids, peptides, and analogs, and triterpenoids subclasses. Some of these components have been previously studied in relation to *P. syringae*. For instance, in the positive polarity we observed prenol lipids or the subclass triterpenoids which are essential for the immune response in plants [38–40]. Additionally, glycerophospholipids are known to be exploited by *Pseudomonas* strains to enhance their interactions with host organisms. [41]. In the negative polarity, *Peltigera* primarily shares carboxylic acids and their derivatives, for which we have not found a direct correlation with the presence of *P. syringae*.

The proportion of shared metabolites in *Peltigera* species is substantially higher in positive polarity (*P. aphthosa* = 75.98%; *P. leucophlebia* = 75.37%; *P. membranacea* = 90.93%) compared to negative polarity (*P. aphthosa* = 7.3%; *P. leucophlebia* = 6.2%; *P. membranacea* = 10.8%). Additionally, *P. membranacea*, which exhibited the lowest number of unique metabolites, showed a higher percentage of shared metabolites with the other *Peltigera* species in both polarities (Table A4).

### The metabolic profiles differentiate lichen species and genera

Based on the previous observation and considering the original hypothesis stated that *Peltigera* may support the growth of *P. syringae* due to the presence of specific metabolites that were absent from other lichen genera, we evaluated the differences between the chemical profiles of *Peltigera* against *Cladonia* and *Stereocaulon*. Similarly to what we observed in the initial analysis of the *Peltigera* species, the chemical profile of the *Peltigera, Cladonia*, and *Stereocaulon* lichens (Figure 2) varies both across genera (positive mode: PERMANOVA F: 4.0841; p= 1.00 × 10^−4^, negative mode: PERMANOVA F: 88.1250; p= 1.00 × 10^−4^) and species (positive mode: PERMANOVA F: 3.1864; p= 1.00 ×10^−4^, negative mode: PERMANOVA F: 335.8548; p= 1.00 ×10^−4^). Clear clustering was observed for both species and genus levels. This observation aligns with previous findings that pointed to secondary metabolites as valuable tools in lichen taxonomy and systematics [29,42,43]. Nevertheless, caution is warranted when attributing differences in metabolites solely to genetic relatedness, as environmental factors can lead to the emergence of chemosyndromes in species [44] or phenomena like homoplasy, which has been studied in lichens, can generate phenotypically cryptic species [45].

**Figure 2.**
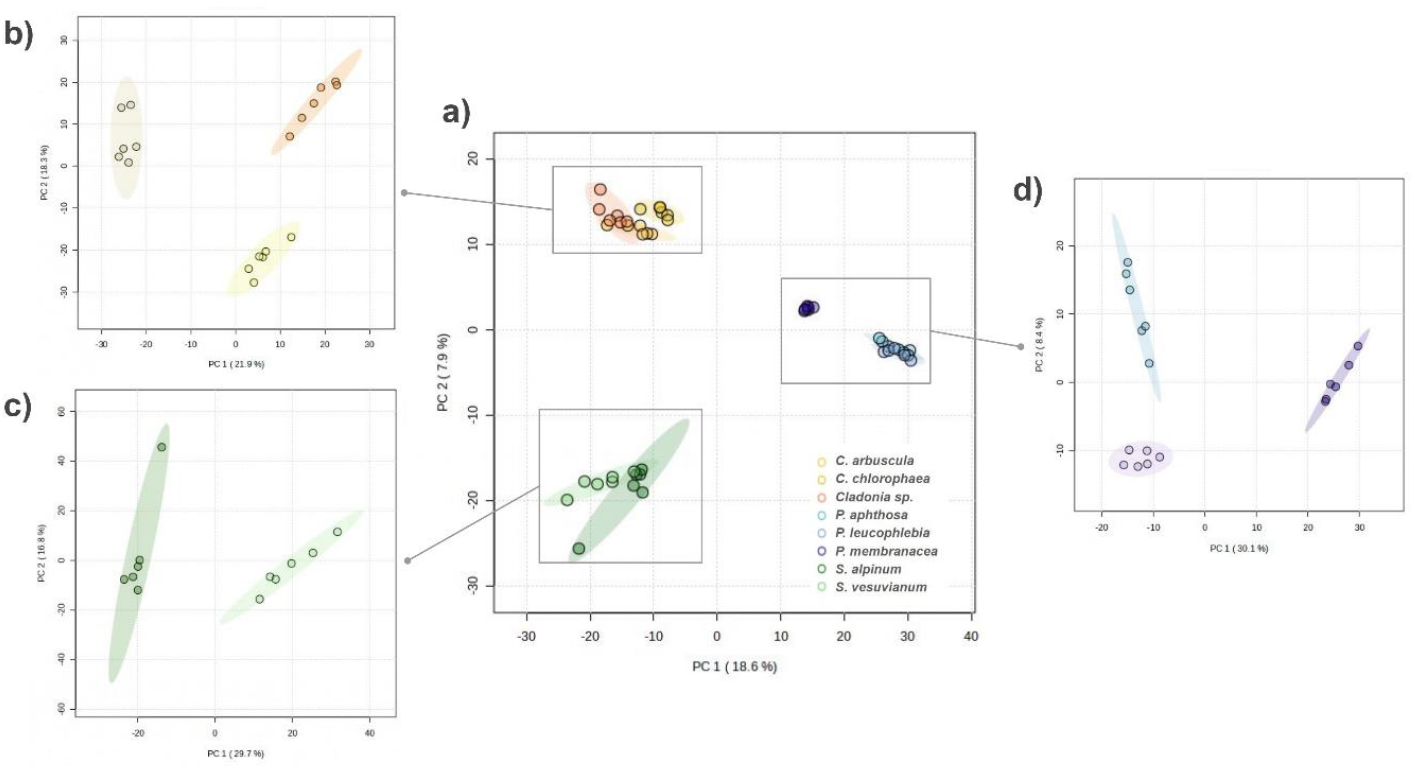
Lichens produce metabolites that cluster by species and by genus. Principal component analysis of the metabolites from ESI positive metabolites isolated from samples utilised in this study a) All samples b) PCA comparing only *Cladonia* samples, c) PCA with a database of only *Stereocaulon* samples, d) PCA comparing only *Peltigera* samples. The image was created by Metaboanalyst.

To determine the main differences between genera, the features detected on each mode (positive or negative) were classified at the superclass and subclass levels using SIRIUS 4 (Figure 3). In the positive mode dataset, lipids (and related compounds) were the more abundant superclass in all the genera aligning with previous studies [46], followed by organoheterocyclic compounds (Figure 3b). *Peltigera* exhibited lower relative abundances of lipid-related molecules in negative polarity compared to other genera. The role of lipids in lichen has been related to the response and adaptation to environmental factors such as temperature, elevation, light, or high levels of sulphur or radiation [47–50]. Notably, the lipids subclass diversity was variable among genera. For example, *Peltigera* lichens had a higher abundance of triterpenoids, while it produced lower amounts of fatty acids, fatty alcohols, and bile acids (Figure A3.a). Organoheterocyclic compounds form a diverse group, encompassing some highly effective fungicides like captan, iprodione, and vinclozolin [51], as well as antiviral and antimicrobial agents such as benzimidazole and pyridines. They also exhibit antioxidant and antiherbicidal properties [52]. Additionally, another common superclass among our samples, phenylpropanoids was less abundant in *Peltigera* and some *Cladonia* than in *Stereocaulon*. Notably, many phenylpropanoids, such as noradrenaline, or *cis N*-*p*-coumaroyloctopamine, have been associated with plant defense against pathogens, including *P. syringae* [53,54].

**Figure 3.**
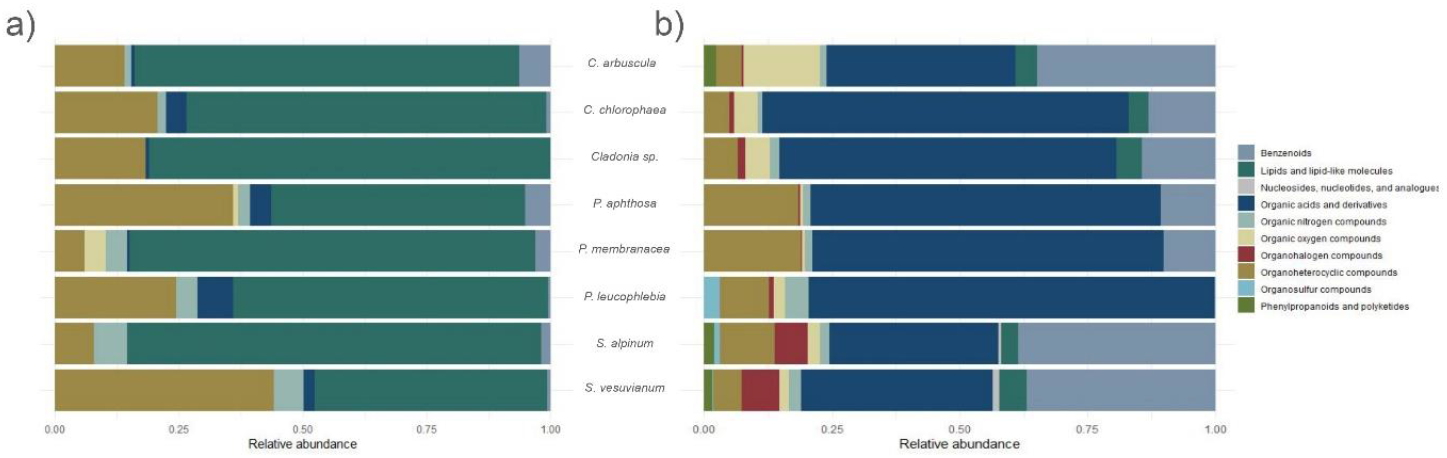
Metabolites grouped by superclass show lipids and organic acids are the most abundant metabolites in the lichens. Identification of compound superclasses from lichens determined by ClassyFier with values expressed in relative abundance. a) Positive polarity b) Negative polarity.

In the negative mode dataset, organic acids and benzenoids were the major superclasses identified (Figure 3b). The levels of organic acids varied among genera, with lower concentrations observed in *Peltigera* compared to the other genera. Specifically, amino acid levels were also lower in *Peltigera* relative to the other two genera, while alpha-keto acids were not detectable in *Peltigera*. Analysis of the benzenoids superclass in the negative mode showed a high abundance in the *Stereocaulon* and *C. arbuscula* metabolic profiles, while they seem to be a minority superclass in *Peltigera*, especially *P. leucophlebia* (Figure 3b). Previous experiments reveal that plants (and microorganisms) generate and release benzenoids in response to stress, leading to think to scientist that their function may be associated with chemical communication and stress protection [55,56]. The subclass benzoic acid and the anisoles showed lower relative abundance in *Peltigera* (Figure A3.b).

### Potential influence of *Peltigera* and non-*Peltigera* metabolites on *P. syringae* presence on them

To test whether *Stereocaulon* and *Cladonia* produce unique metabolites that might be inhibitors or deleterious for *P. syringae*, we examined whether there were unique or more abundant metabolites present in non-*Peltigera* genera. To do this, we examined the 333 metabolites identified from a PLS-DA model as class discriminant features (VIP>1.5) that distinguish *Peltigera* and non-*Peltigera* lichen genera (*Cladonia* and *Stereocaulon*). Among the selected features, 28 were found in both *Peltigera* and non-*Peltigera*, 115 were exclusive to *Peltigera*, and the remaining 190 were exclusive to non-*Peltigera* genera. Among the VIPs identified, 77% were lipids, the most prevalent chemical family, followed by organic acids and their derivatives (Figure 4; Table A5).

**Figure 4.**
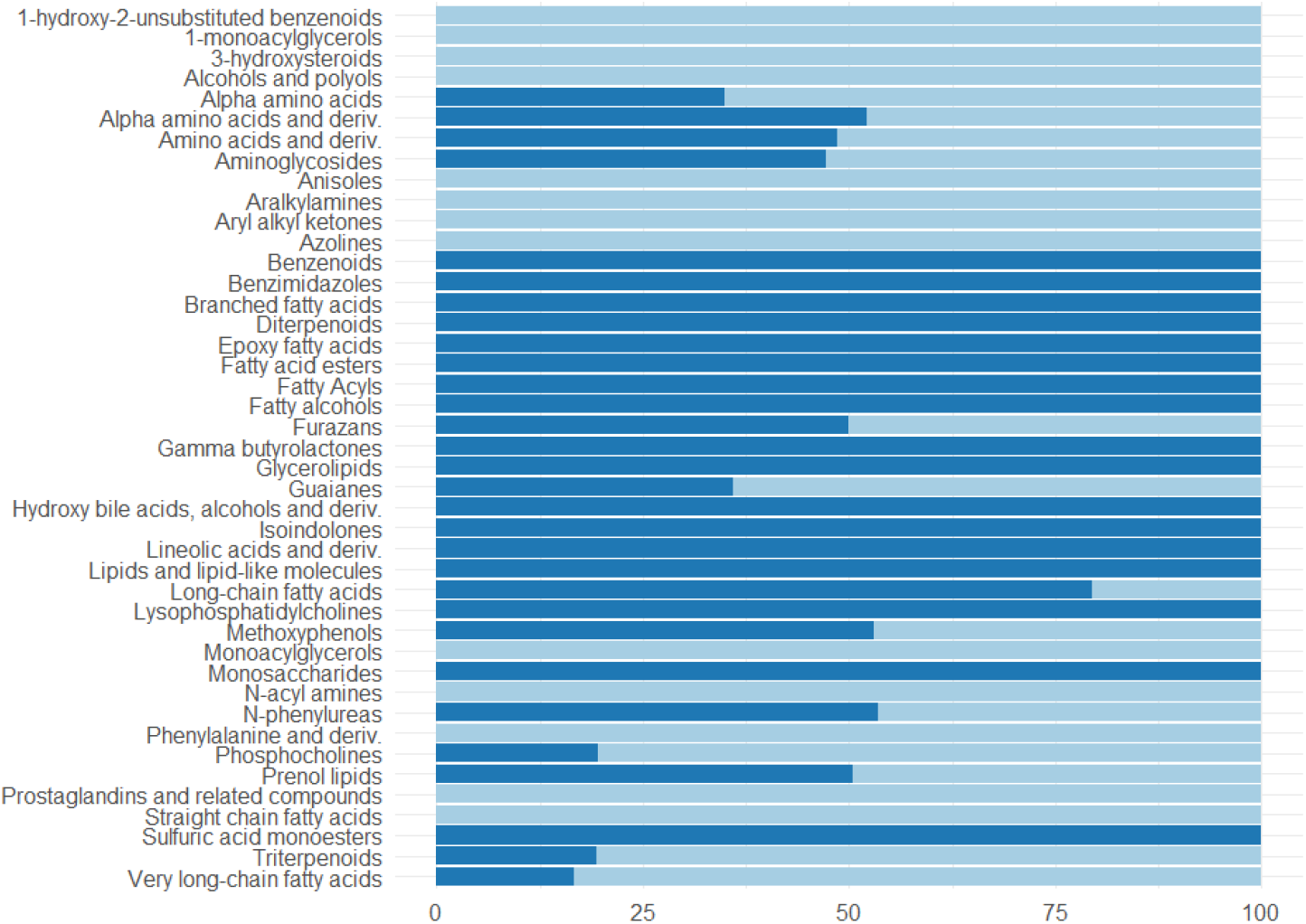
VIP compounds abundance differenciating Peltigera and non-Peltigera samples annotated as most specific class. Light blue represents non-Peltigera while dark blue is Peltigera.

Remarkably, several metabolites with known antimicrobial properties were identified only in *Peltigera*. This is the case of aminoglycosides a diverse group of antibiotics which some of them affect a wide range of bacteria including *P. syringae* and some of their antagonist as *B. subtilis, P. aeruginosa, Enterobacter*, and *Serratia* sp [57,58]. Among the predominant compounds in *Peltigera* we had pyridinones which has antimicrobial effect against some gram-negative bacteria as *P. aeruginosa* [59,60]. On the other hand, some metabolites with antimicrobial properties were only identified in non-*Peltigera* such as cyclic depsipeptides or azolines which inhibit *P. aeruginosa, B. subtilis, Staphylococcus aureus* among others. In the negative mode, the potent antimicrobial usnic acid [20] was almost absent on the *Peltigera* analysed samples. Usnic acid strongly inhibits some of the *P. syringae* competitors as *B. subtilis* and *S. aureus* [61]. However, usnic acid is not active against gram negative bacteria [62].

Some metabolites predominantly present in *Cladonia* and *Stereocaulon* might inhibit the presence of *P. syringae* in their thallus by other methods. This is the case of ubiquinones identified as regulators of basal resistance in plants against *P. syringae* and stress mediated by reactive oxygen species [63]. Also, salicylic acid was only isolated from non-*Peltigera* genera, and this has been observed to be produced by different plant species after *P. syringae* infection [64–66] with a key role in plant resistance [67]. Choline levels are notably higher in the non-*Peltigera* samples and has been reported to enhance plant growth and as well as protection from plant pathogens including *P. syringae* and *B. subtilis* [68]. Contrary, we identified compounds only in *Peltigera* classified as benzimidazoles which are a set of compounds that play a role in *P. syringae*-plant interactions, not inhibiting *P. syringae* growth, but providing protection to the plant [69]. Also, triterpenoids mainly found in *Peltigera* are involved in protection of plants against pathogens [40], which might also be the role in lichens. Prostaglandins absent in *Peltigera* are related with defence mechanisms in microalgae [70]. *Peltigera* exhibited an absence of xanthines in both polarities but was found in the other two genera. The enzyme xanthine oxidoreductase exhibits superoxide-producing activity, effective against several pathogens but not with *P. syringae* [71].

Additionally, the non-detection of anthracene in *Peltigera* (Table A6), used by species like *B. subtilis, B. velezensis*, and *P. aeruginosa* for nutrition, implies a substrate competition that could inhibit *P. syringae’s* presence [72–75]. Based on the observations of these metabolic analyses, the data suggest that there could be specific metabolites supporting *Peltigera* growth or inhibiting it in *Stereocaulon* and *Cladonia*.

### All lichen tested support *P. syringae* growth

One potential reason that *P. syringae* is only isolated from *Peltigera* is that this lichen produced something that other lichen species do not. To test this, growth assays using lichen extracts were conducted on a range of *P. syringae* isolates and compared against growth in rich medium (KB); *B. velezensis* was used as an outgroup control. Given the number of strains and growth substrates being used, four growth trials were carried out where two lichen species were used in each trial. The growth of strains in each trial was compared as pairs. Overall, growth in KB was generally 3-4 times higher (reaching OD_600_ values of >1.28) than growth in the M9 (reaching OD_600_ values of 0.4111 in M9 + lichen extract; no strains were able to grow in M9 without lichen extract). All *P. syringae* isolates (except CFBP1392 in *Stereocaulon*) were able to grow in M9 supplemented with lichen extracts (Figure 5). Variations in growth were observed between *Cladonia* sp. and *C. arbuscula* extracts, with all strains growing best in the medium supplemented with *Cladonia* sp. (*Cladonia* sp. maximum OD_600_: 0.12-0.2; *C. arbuscula* maximum OD_600_: 0.005-0.06). Also, bacterial growth comparison among *P. syringae* phylogroups calculated with all samples pooled together showed a statistically higher growth in PG13 compared to PG2 in lichens, an effect not observed in enriched control media (KB), which supports both PG2 and PG13 equally and to a lesser extent, PG10 (Figure A7).

**Figure 5.**
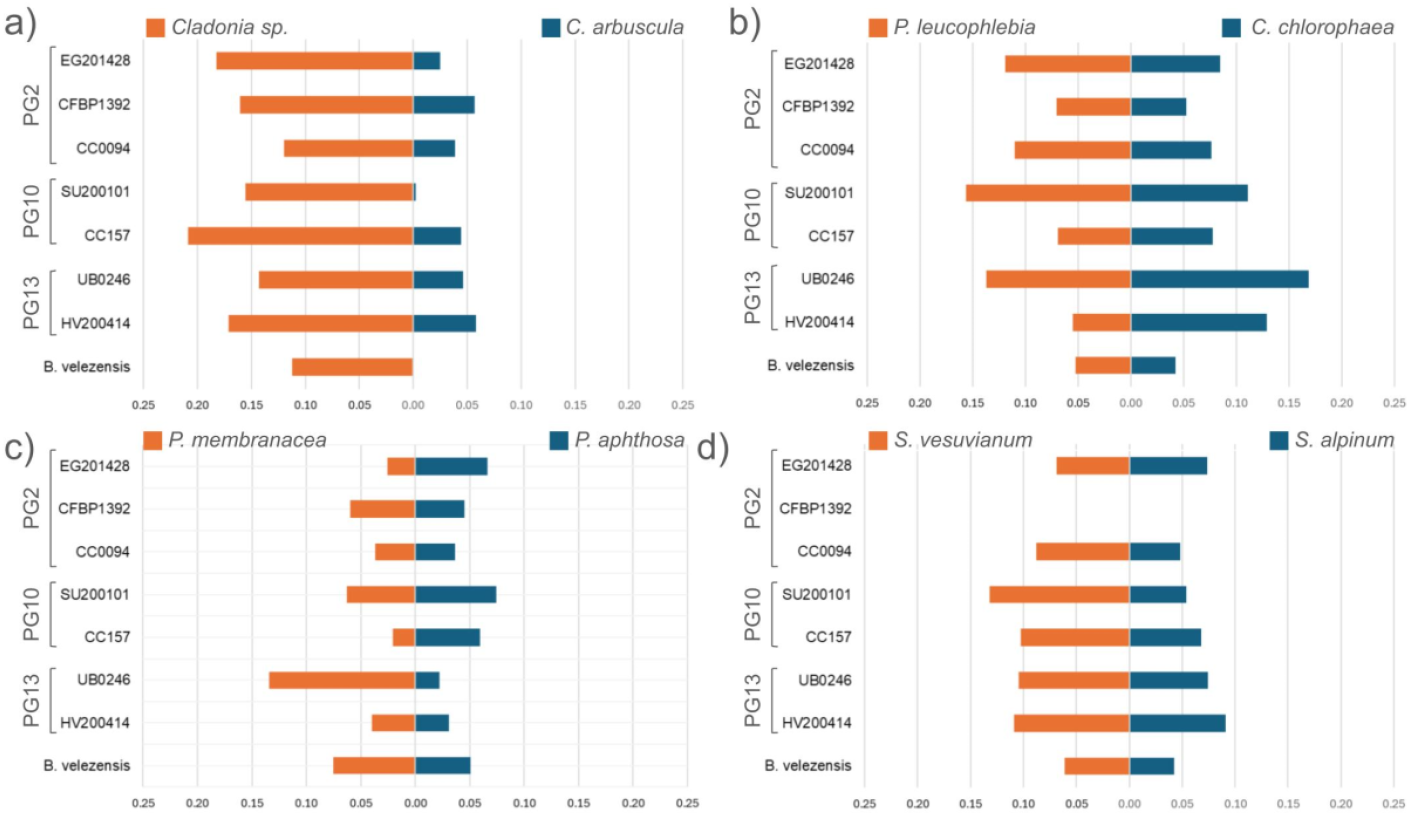
*P. syringae* can utilise lichen extracts for growth. The bacterial culture density (OD_600_) was observed after 47 hrs incubation in M9 minimal medium containing different lichen extracts. Values are presented by pairs tested in the same trials and the values represent the average of three replicates. *a) C. arbuscula* and *Cladonia sp*.; *b) C. chlorophaea* and *P. leucophlebia; c) P. apthosa* and *P. membranacea; d) S. alpinum* and *S. vesuvianum*.

One possible reason for why *P. syringae* is able to colonise *Peltigera* is that it can tolerate the lichen environment compared to other lichens. To test this, the antimicrobial properties of lichen extracts from each studied species was used against different *P. syringae* bacteria isolated from *Peltigera* lichen and other strains isolated from plants of snow from three different countries as well as *B. velezenzis*. Notably, clear halos indicating bacterial growth inhibition were observed only in *B. velezensis* and not *P. syringae* strains (Figure 6). This effect was only noticed when using lichen extracts from *Cladonia* and *Stereocaulon* species (Figure 6a), but no inhibitory effect was observed when using the *Peltigera* extract, indicating that this lichen may be a more microorganism-permissive holobiont. This indicates the host-specificty effect is unlikely to be due to toxic products being produced by *Cladonia* and *Stereocaulon*. Interestingly, all lichens caused the appearance of a hazy halo for some of the *P. syringae* strains, appearing ‘wetter’ than the rest of the plate (Figure 6). This might indicates the production of a surfactant by the bacterium. This phenomenon previously reported as a result of surfactin production by *P. syringae* [76], appears to vary between *P. syringae* strains, with a wider halo observed in PG13, followed by PG10, and finally PG02 (Figure 6b).

**Figure 6.**
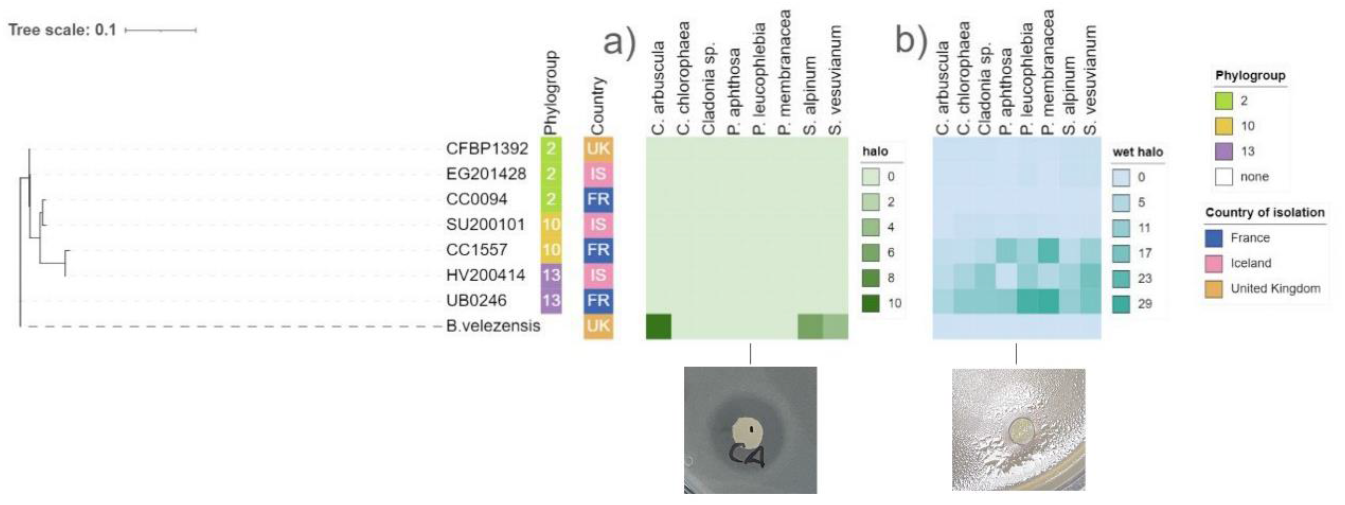
*Cladonia* and *Stereocaulon* inhibit *B. velezensis* but trigger some *P. syringae* to produce a surfactant-like ‘wet’ halo. Selected *P. syringae* strains from different phylogroups are shown as well as B. velezensis. The diameter of halos (in millimetres) is colour-coded a) inhibition halo, b) ‘wet’ halo.

## Conclusion

In this study, we explored the metabolic profiles of various lichens belonging to the genera *Cladonia, Stereocaulon*, and *Peltigera*. Our aim was to identify distinctive or commonly occurring metabolites within these lichens, with the intention of understanding how specific metabolites or their absence might contribute to the exclusive isolation of *P. syringae* from *Peltigera*. We should be cautious when interpreting the metabolites identified in different lichens in this study, as the lichens were cleaned but not sterilised. It’s possible that metabolites from other organisms present on the lichen surface, rather than within the lichen thallus itself, were also observed. Additionally, factors such as sampling location, lichen age, or varying health status of the lichens might explain why two of the samples were considered as *Cladonia* sp. instead of *C. chlorophaea*, due to differences not only in the metabolic profile but also in the kinetic analyses.

Our kinetic and inhibition analysis did not show *Peltigera* supporting *P. syringae* growth to a greater extent than the other lichens, as would be expected from the isolation efforts described in Ramírez et al. (2023) where no *P. syringae* was isolated from non-*Peltigera* lichens. Notably, however, for *C. arbuscula* there was a clear difference of this lichen as nutrient source for *P. syringae* compared with the rest of the lichen analysed in the study. Additionally, it raises the possibility that *Peltigera* actively supports the presence of *P. syringae* while the lichen is alive, adding complexity to the study due to the challenge of maintaining lichens in laboratory conditions. Another factor might be the structural differences in the cell wall among lichens, which could facilitate *P. syringae*’s entrance into the thallus [77]. A plausible explanation could be that *Peltigera* lichens retain more water compared to the other two genera analyzed in this study. However, it is important to note that all these lichens are poikilohydric and dry out quickly (Table A1). This may be reasonable given the strong association of *P. syringae* with water, as it is commonly found in various surface water masses.

While our findings did not pinpoint a specific metabolite as the cause of *P. syringae’s* selective behaviour, we identified certain metabolites that could potentially play a role. We hypothesise that this difference might be produced by a combination of metabolites or the loss of some metabolite that might be important during the extraction or other part of the process. We highlight the complexity of the lichen metabolic profile being significantly higher in *Cladonia* and *Stereocaulon* than in *Peltigera*. The extensive diversity of compounds presents a remarkable chemical complexity for organisms interacting with lichens. This could explain why a less diverse compound profile provided by *Peltigera* might be more favorable to *P. syringae*.

## Supporting information

Description of the samples analyse with the species that belong, place of isolation, water content, sampling data, and symbionts.

Abundance of subclasses within the predominant superclass in each polarity. a) "lipid-like molecules" from positive polarity

Abundance of subclasses within the predominant superclass in each polarity. b) organic acid and derivates from negative polarity

List of shared metabolites level between Peltigera samples annotated to class and subclass in both polarities and shared metabolites abundance

List of VIP>1.5 Peltigera vs non-Peltigera comparison

List of features from all samples with annotations at 60% confidence level or higher from both ESI modes

Bacterial growth of P. syringae strains and B. velezensis color by PG and with line type by isolation place

## Appendix

Table A1. **Description of the samples analyse with the species that belong, place of isolation, water content, sampling data, and symbionts**. The missing information is marked with an “X”.

**Table A2.**
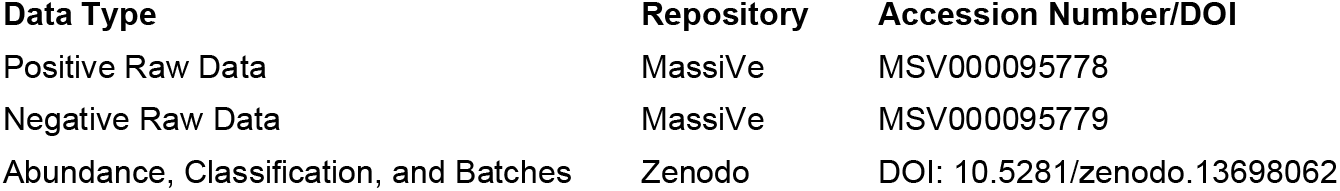
MzMine batch files, metabolite abundance and clasification of metabolites from both polarities:

Figure A3. **Abundance of subclasses within the predominant superclass in each polarity**. The abundance of the subclasses was converted to Log10 to enhance the clarity of representation for all compounds) “lipid-like molecules” from positive polarity b) organic acid and derivates from negative polarity

Table A4. **List of shared metabolites level between *Peltigera* samples annotated to class and subclass in both polarities and shared metabolites abundance**

Table A5. **List of VIP>1.5 *Peltigera* vs non-*Peltigera* comparison**. Abundance is expressed in arbitrary units (AU).

Table A6. **List of features from all samples with annotations at 60% confidence level or higher from both ESI modes**

Figure A7. **Bacterial growth of *P. syringae* strains and *B. velezensis* color by PG and with line type by isolation place** (“IS”-Iceland or “O” – other). Separate graphs represent different media types, with numbers indicating the trial number.

## Acknowledgments

This work was funded by the Rannís Icelandic Research Fund under project number 1908-0151.

