## Supplementary figures and images for "Exploring the Exclusive Isolation of *Pseudomonas syringae* in *Peltigera* Lichens via metabolite analysis and growth assays"

### Abundance of subclasses within the predominant superclass in each polarity. a) "lipid-like molecules" from positive polarity

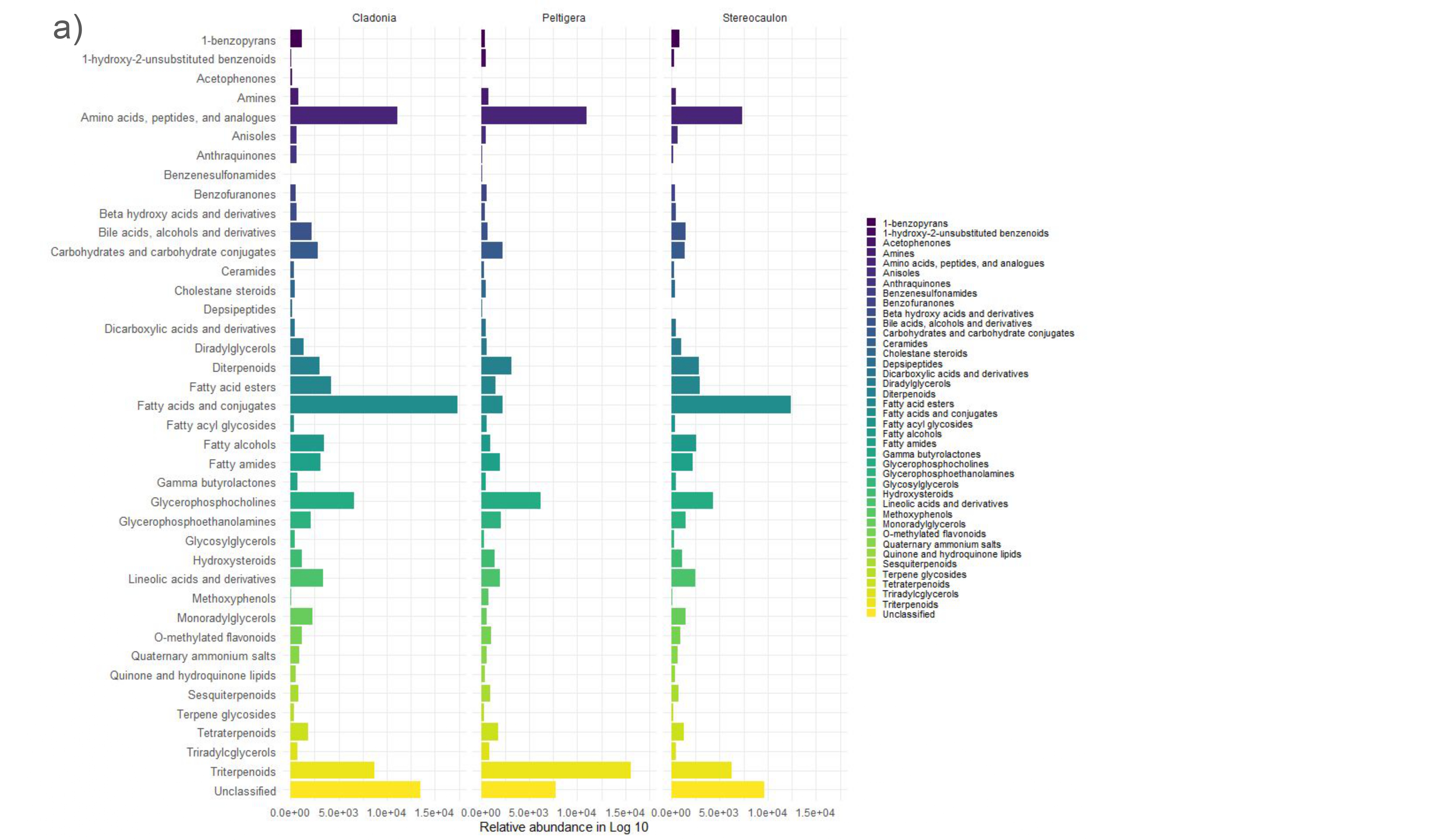

### Abundance of subclasses within the predominant superclass in each polarity. b) organic acid and derivates from negative polarity

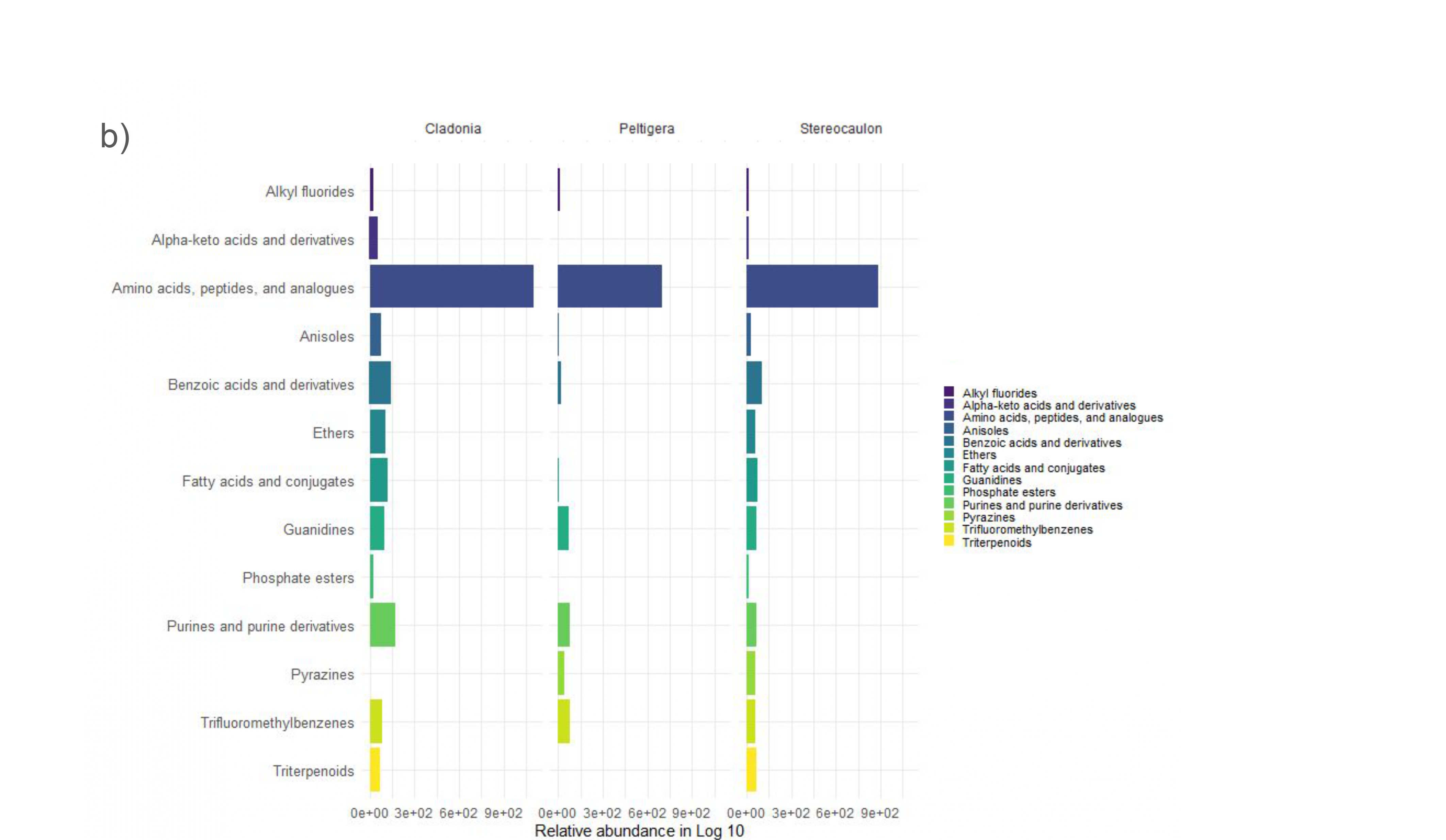

### Bacterial growth of P. syringae strains and B. velezensis color by PG and with line type by isolation place

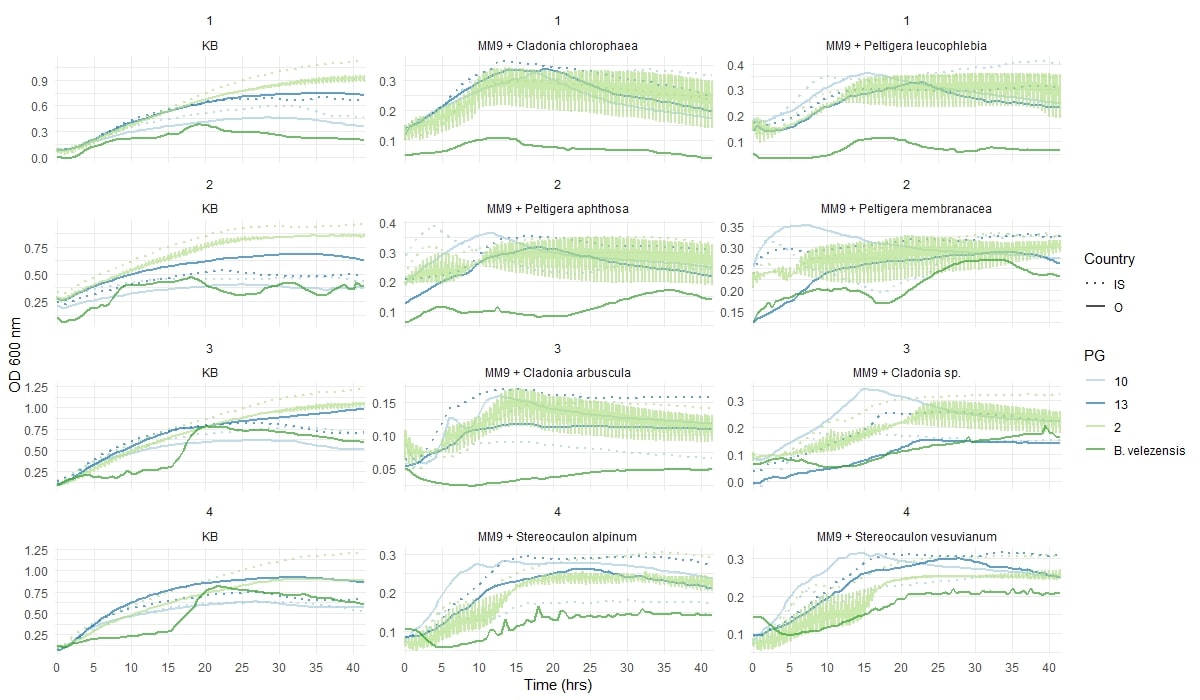
